# Characterization, enrichment, and computational modeling of cross-linked actin networks in trabecular meshwork cells

**DOI:** 10.1101/2024.08.21.608970

**Authors:** Haiyan Li, Devon H. Harvey, Jiannong Dai, Steven P. Swingle, Anthony M Compton, Chenna Kesavulu Sugali, Kamesh Dhamodaran, Jing Yao, Tsai-Yu Lin, Todd Sulchek, Taeyoon Kim, C. Ross Ethier, Weiming Mao

## Abstract

**Purpose:** Cross-linked actin networks (CLANs) are prevalent in the glaucomatous trabecular meshwork (TM), yet their role in ocular hypertension remains unclear. We used a human TM cell line that spontaneously forms fluorescently-labeled CLANs (GTM3L) to explore the origin of CLANs, developed techniques to increase CLAN incidence in GMT3L cells, and computationally studied the biomechanical properties of CLAN-containing cells.

**Methods:** GTM3L cells were fluorescently sorted for viral copy number analysis. CLAN incidence was increased by (i) differential sorting of cells by adhesion, (ii) cell deswelling, and (iii) cell selection based on cell stiffness. GTM3L cells were also cultured on glass or soft hydrogel to determine substrate stiffness effects on CLAN incidence. Computational models were constructed to mimic and study the biomechanical properties of CLANs.

**Results:** All GTM3L cells had an average of 1 viral copy per cell. LifeAct-GFP expression level did not affect CLAN incidence rate, but CLAN rate was increased from ∼0.28% to ∼50% by a combination of adhesion selection, cell deswelling, and cell stiffness-based sorting. Further, GTM3L cells formed more CLANs on a stiff vs. a soft substrate. Computational modeling predicted that CLANs contribute to higher cell stiffness, including increased resistance of the nucleus to tensile stress when CLANs are physically linked to the nucleus.

**Conclusions:** It is possible to greatly enhance CLAN incidence in GTM3L cells. CLANs are mechanosensitive structures that affect cell biomechanical properties. Further research is needed to determine the effect of CLANs on TM biomechanics and mechanobiology as well as the etiology of CLAN formation in the TM.

## Introduction

Glaucoma, a major cause of blindness, is a common optic neuropathy in which retinal ganglion cell dysfunction and damage result in characteristic patterns of visual field loss. Ocular hypertension (OHT) is a major risk factor for glaucoma; moreover, it is the only treatable risk factor. Ocular hypertension in primary open-angle glaucoma (POAG), the most common form of glaucoma, is due to elevated aqueous humor outflow resistance, which in turn is most frequently caused by pathological changes in the trabecular meshwork (TM) and inner wall of Schlemm’s canal (SC), as reviewed in (Stamer and Clark 2017).^1^ Pathological findings in the TM associated with OHT include: loss of TM cells,^2–4^ compromised TM cell function, excessive extracellular matrix (ECM) deposition, and increased TM^5–7^ and ECM stiffness^6, 8^. Despite these observations, the fundamental causes of OHT remain unknown.

In addition to the pathological changes described above, it is known that cross-linked actin networks (CLANs) are associated with OHT: they occur more frequently in TM cells from glaucomatous eyes and can be induced by agents known to cause OHT, such as TGF-β2 and dexamethasone (DEX).^9–16^ These studies strongly suggest a functional link between CLANs and OHT. CLANs consist of interconnected filamentous (F)-actin (“hub and spoke” morphology), appearing as web-like (in 2D images) or spherical (in 3D images) intracellular structures.^11, 12^ We and others have shown that CLANs colocalize with multiple proteins, including PIP2, syndecan, α-actinin, filamin, PDLIM1, caldesmon, calponin, and tropomyosin.^13, 17^

Despite the significant associations between CLANs and OHT/glaucoma, much remains unknown about CLANs, including their impact on cellular functions, ECM production/remodeling, mechanotransduction, and most importantly, intraocular pressure (IOP). These knowledge gaps are due in part to experimental barriers. For example, it is difficult to induce and visualize CLANs, and thus the reproducibility of CLAN studies has been challenging. Prior to our recent paper, most previous CLAN-related research used primary human TM (pHTM) cells together with glucocorticoid, TGF-β2, or integrin activation, with the exception of some studies using bovine TM cells.^11–14, 16–20^ These approaches have drawbacks; for example, CLAN induction rate varies significantly between cell strains and even between batches from the same cell strain, hindering reproducibility and rigor. Moreover, the use of primary cells is inherently limited by passage numbers,^21^ making it challenging to conduct experiments that require significant numbers of cells.

We recently described a unique TM cell line, GTM3-LifeAct-GFP (GTM3L). The GTM3L cell line was derived from the widely-used GTM3 cell line, established about 20 years ago by immortalizing glaucomatous pHTM cells.^22, 23^ GTM3 cells share many features with pHTM cells, including phagocytic capability and DEX-inducible myocilin expression.^24–26^ We serendipitously discovered the GTM3L subline, which spontaneously forms fluorescently labelled CLANs suitable for live imaging.^22^ Using these cells, we showed that CLANs make TM cells stiffer, less dynamic, and more resistant to latrunculin-B (an actin polymerization inhibitor).^22^

Interestingly, not all GTM3L cells form CLANs.^22^ To further study CLANs using GTM3L cells, it is important to be able to enrich CLAN+ cells beyond their spontaneous low incidence rate (defined as the number of CLAN+ cells divided by the total number of cells) reported in GTM3L cells ^22^. In this study, we explored the potential origin of CLANs, combined several methods to increase CLAN incidence rate in GMT3L cells with validation of their effects on cell stiffness, and developed a computational model to study the biomechanical impact of CLANs on cells.

## Methods

### Cell culture

GTM3L cells were cultured in low-glucose Dulbecco’s Modified Eagle’s Medium (DMEM; Gibco, Thermo Fisher Scientific, Waltham, MA, USA) or Opti-MEM (Thermo Fisher Scientific) containing 10% fetal bovine serum (FBS; HyClone, Thermo Fisher Scientific) and 1% penicillin/streptomycin/glutamine (PSG; Gibco/Thermo Fisher Scientific), and maintained at 37°C in a humidified atmosphere with 5% CO_2_. Fresh media was supplied every 2-3 days.

### Fluorescence-activated sorting and Lentiviral copy number analysis

About 2 x 10^7^ GTM3L cells were sorted into 3 groups with high, medium, and low GFP intensity using a BD FACSAria™ III Cell Sorter (BD, Franklin Lakes, NJ) at the core facility at Indiana University (Supplemental Figure 1). These cells were cultured, and DNA was isolated using a Nucelospin kit (Macherey-Nael) for viral copy number analysis. The copy number of the lentiviral vector sequence in the Woodchuck Hepatitis Virus Posttranscriptional Regulatory Element (WPRE) was normalized to the copy number of the host cell’s ApoB genomic sequence using droplet digital PCR (Bio-rad, Hercules, CA). The primers, PCR conditions, and reagents are listed in Supplemental Materials.

### Hydrogel preparation

The hydrogel precursor gelatin methacryloyl (GelMA [6% w/v final concentration], Advanced BioMatrix, Carlsbad, CA, USA) was mixed with lithium phenyl-2,4,6-trimethylbenzoylphosphinate (LAP, 0.075% w/v final concentration) photoinitiator (Sigma-Aldrich, Saint Louis, MO). Thirty microliters of the hydrogel solution were pipetted onto Surfasil-coated (Thermo Fisher Scientific) 18 × 18-mm square glass coverslips followed by placing 12-mm round silanized glass coverslips on top to facilitate even spreading of the polymer solution. Hydrogels were crosslinked by exposure to UV light (CL-3000 UV Crosslinker; Analytik Jena, Germany) at 1J/cm^2^. The hydrogel-adhered coverslips were removed with fine-tipped tweezers and placed hydrogel-side facing up in 24-well culture plates (Corning; Thermo Fisher Scientific).

### Osmotic deswelling of GTM3L cells

GTM3L cells were seeded at 3 × 10^4^ cells/cm^2^ atop either glass coverslips or soft hydrogels and cultured in DMEM with 10% FBS and 1% PSG overnight. Then, GTM3L cells were cultured in DMEM with 1% FBS and 1% PSG and exposed to 2% Polyethylene glycol 300 (PEG300; Sigma) for 2 or 5 days.

### Live cell imaging

Live imaging was used to determine how long CLANs persisted after PEG300 removal. CLANs were defined as “F-actin–containing cytoskeletal structures with at least one triangulated actin arrangement consisting of actin spokes and at least three identifiable hubs.”^18^

GTM3L cells were seeded in either a 35 mm dish with a 0.17 mm thick glass bottom (World Precision Instruments, Sarasota, FL) or on a glass chip (chip ID: CGF400800F; ARRALYZE - part of LPKF Group, Garbsen, Germany) which was mounted on a 35mm plastic dish with a 22 mm square cutout in the center. The glass chip contained 512 microwells of 400 µm diameter and 162 microwells of 800 µm diameter. The bottom of the microwell was about 175 um thick, and the height of the wall of the microwell was about 475 µm. The behavior of cells on both substrates was similar and the data we present combines experiments from both substrates. The cells were cultured in serum-free OptiMEM supplemented with 1% glutamine/penicillin/streptomycin. On treatment day 0, the cells were treated with 2% PEG300 (Catalog # 202371, Sigma Aldrich). The cells then were monitored daily for the formation of CLANs, which most often formed between treatment days 4-6. Once CLANs were identified, treatment was withdrawn by gently replacing the medium containing 3% PEG300 with fresh culture media without PEG300. Cells were immediately transferred to and were maintained in a stage top incubator (Tokai, Shizuoka-ken, Japan) secured on the stage of a Nikon Eclipse Ti2 inverted microscope (Nikon, Melville, NY) set at 5% CO_2_ and 37°C for the duration of live imaging. Time lapse images were immediately captured using a 40x objective S plan fluor ELWD objective (Nikon). Further images were captured every 30 seconds for the first 2 hours and then at every minute for an additional 4 hours.

### Image analysis

To determine the persistence time of CLANs after PEG300 withdrawal, five researchers individually analyzed the captured images. The time at which CLANs were no longer visible was recorded, and the average time over the 5 observers was taken as the persistence time of CLANs after PEG300 withdrawal. For this part of the study, our definition of visible CLANs was identical to that described above in “Live cell imaging”.

### Cell sorting based on cell-substrate adherence

GTM3L cells were plated in T75 flasks and grown to confluence in DMEM with 10% FBS and 1% PSG. 2 ml of 0.25% trypsin was added and incubated for 2 min at 37°C, and non-adherent cells and media were discarded. The remaining adherent cells were then released by adding a further 2 ml of 0.25% trypsin for 30 s, adding media to neutralize the trypsin, centrifugation for 10 min at 1000g, and resuspending the pellet in DMEM with 10% FBS and 1% PSG. The cells were then seeded on glass coverslips for testing osmotic deswelling-induced CLAN formation.

### Cell sorting based on cell stiffness

GTM3L cells were selected by stiffness according to an established protocol.^27–31^ In brief, 2 ml of a GTM3L cell suspension (1.8 x 10^6^ cells/ml) were passed through a microfluidic device with a narrow channel designed with 14, 7 µm constrictions angled at 30° that direct the trajectory of cells, and thus fractionate cells, depending on their stiffness. This device was made from PDMS (Sylgard 184; Thermo Fisher Scientific) cured on an SU-8 photoresist mold etched using standard photolithography procedures. The day before sorting, devices were passivated with 1% Pluronic F-68 solution (Thermo Fisher Scientific). Cells suspended in flow buffer (culture media with 20% Percoll to provide neutral buoyancy to cells and 0.4 mg/ml DNAse I to avoid DNA-induced blockage if cells lyse) were flowed into the device at 3-7 μl/min and were collected from the device outlets (each corresponding to a different stiffness), while control samples were taken from unperfused cells. Samples were spun down and resuspended in 150 μl of fresh culture media, and the suspended cells were plated and expanded.

### Atomic force microscopy

An MFD-3D AFM (Asylum Research, Santa Barbara, CA, USA) was used to make cell stiffness measurements using silicon nitride cantilevers with an attached borosilicate sphere (diameter = 10 μm; nominal spring constant = 0.1 N/m; Novascan Technologies, Inc., Ames, IA, USA). Cantilevers were calibrated by measuring the thermally induced motion of the unloaded cantilever before measurements. The indentation depth was limited to 400 nm to avoid substrate effects and the tip velocity was adjusted to 800 nm/s to avoid viscous effects.^32^ Five measurements/cell were conducted, and at least 5 cells were measured/group. For hydrogel stiffness measurement, a force map covering a 40 x 40 μm area (5 x 5 grid of points) was measured. Data from AFM measurements were fitted to the Hertz model to calculate the effective Young’s Modulus of the cells, assuming a Poisson’s ratio of 0.5.

### Immunostaining

GTM3L cells were fixed with 4% paraformaldehyde (Electron Microscopy Sciences, Hatfield, PA, USA) at room temperature for 20 min, permeabilized with 0.5% Triton™ X-100 (Thermo Fisher Scientific) and incubated with Phalloidin-iFluor or 594 (Cell Signaling Technology, Danvers, MA, USA)/DAPI according to the manufacturer’s instructions. Coverslips were mounted with ProLong™ Gold Antifade (Invitrogen) on Superfrost™ microscope slides (Theromo Fisher Scientific), and fluorescent images were acquired with a Leica DM6 B upright microscope system. Images were captured from at least 10 fields per group, which corresponds to over a thousand cells. The images were subsequently analyzed to determine the percentage of CLAN+ cells, calculated as the ratio of the number of CLAN+ cells to the total number of cells.

### Computational modeling

To simulate CLAN-like networks, we employed our agent-based model with F-actin and actin cross-linking protein (ACP) simplified via cylindrical segments.^33–36^ (Supplemental Figure 2) A cell nucleus was included in some simulations, represented as a triangulated mesh. The positions of all points defining the cylindrical segments and the triangulated mesh were updated in each time step using the Langevin equation and the forward Euler integration scheme. For all elements, extensional, bending, and repulsive forces were considered as deterministic forces. For the nucleus, forces enforcing conservation of volume and surface area were also considered. F-actins were assembled via nucleation with specific orientations and polymerization, but they did not undergo depolymerization. Actin cross-linking proteins interconnected pairs of F-actins to form functional cross-linking points.

Via the self-assembly process of F-actin and ACP, CLAN-like networks were created. In simulations without the cell nucleus, the CLAN-like network was formed in a thin rectangular domain (10 × 8.66 × 0.1 µm) with periodic boundary conditions in the x and y directions. The actin concentration (*C*_A_) was 100 µM, and the molar ratios of ACPs (*R*_ACP_ = *C*_ACP_/*C*_A_) was 0.1. After network assembly, the periodic boundary condition was disabled in the y direction. Then, F-actins crossing the two boundaries normal to the y direction were severed, and their ends were clamped to the boundaries. For bulk rheology measurement, the +y boundary was displaced in either the +x, +y, or -y directions at a constant rate to apply shear, tensile, or compressive strain to the network, respectively, while the -y boundary was fixed. The maximum strain was 0.05, and the strain rate was 0.001 s^-1^. At each strain level, stress was calculated by summing the component of forces acting on the ends of all F-actins clamped on the +y boundary and then dividing the sum by the boundary area. For shear stress, the x component of forces was used, but for tensile and compressive stresses, the y component was used. Simulations with the cell nucleus were performed in a larger rectangular domain with or without the CLAN-like network. The CLAN-like network was created along the nucleus surface as explained earlier, in a three-dimensional rectangular domain (10×10×5.1 µm). Note that the initial z dimension of the domain is close to the diameter of the nucleus, 5 µm. The actin structure was created right above the nucleus (i.e., within a space defined by a radial distance from the nucleus center between *r*_N_ and 1.1×*r*_N_ where *r*_N_ is a nucleus radius). During the actin assembly, the nucleus was frozen without a change in its spherical shape. *C*_A_ was 60 μM, which was calculated using the space for actin assembly. *R*_ACP_ was 0.1. After network assembly, compressive strain was applied to the +y boundary, whereas the -y boundary was fixed. The maximum compressive strain was -0.2 (i.e., 20% decrease in the z dimension of the domain), and the strain rate is -0.01 s^-1^. In each strain level, a total resistant force exerted by the actin structure and the nucleus was measured, and then stress was calculated by dividing the total force by the contact area between the nucleus and the boundary. Further details of the agent-based model and parameters used in simulations are given in Supplemental Materials.

### Statistical analysis

GraphPad Prism software v10.2.3 (GraphPad Software, La Jolla, CA, USA) was used for all analyses. All data sets were tested for normality using the Shapiro-Wilk test and were confirmed to meet the normality criteria. The significance level was set at p<0.05. Comparisons between groups were assessed by t-tests and one-way analysis of variance (ANOVA) with Tukey’s multiple comparisons post hoc tests.

## Results

### GTM3L cells had 1 viral insertion per cell on average

Although the original GTM3 cells were monoclonal, after lentiviral (pLenti-LifeAct-EGFP-BlastR)

^37^ transduction and antibiotic selection, GTM3L cells were established to be polyclonal and observed to have variable GFP intensities. We hypothesized this variation was related to different expression levels of the LifeAct-GFP fusion protein and conducted studies to text this hypothesis.

We sorted GTM3L cells into 3 groups with high, medium, and low GFP intensity, respectively (Supplemental Figure 1). These cells were cultured, and DNA was isolated for viral copy number analysis. We found that naïve GTM3 cells had negligible viral copy number per cell, with readings at background levels. In contrast, all 3 groups of GTM3L cells had 1 viral copy per cell on average (**Table 1**), regardless of their GFP intensity, suggesting that the number of viral copy/insertions per cell was not correlated with LifeAct-GFP expression levels.

**Table 1.** Average viral copy number per GTM3L cell.

| Cells | Average viral copy number per cell |
| --- | --- |
| Naïve GTM3 (negative control) | 9.0439x10<sup>-05</sup> |
| GTM3L high GFP intensity | 1.0999 |
| GTM3L medium GFP intensity | 1.1299 |
| GTM3L low GFP intensity | 1.1320 |

We found that CLAN formation rate was very low among all 3 groups of GTM3L cells described previously (high, medium, and low GFP intensities), and was similar to unsorted GTM3L cells (data not shown). Since CLANs were relatively easy to identify in GTM3L cells with medium and high GFP expression, these two groups of cells were used in subsequent studies.

### Enrichment of CLAN-forming cells based on cell-substrate adhesion, osmotic deswelling and cell stiffness

#### Enrichment of CLAN+ cells based on cell-substrate adhesion

It has been observed that interactions between cells and substrates can enhance the formation of CLANs in TM cells.^17, 19, 38^ This insight led us to employ an adhesion-based selection strategy to augment the proportion of CLAN+ cells. We noted that after 9 cycles of selecting GTM3L cells with stronger substrate attachment, there was a remarkable increase in the frequency of CLAN+ cells, rising from a low incidence of 0.28 ± 0.42% to 4.16 ± 2.95% (**Figure 1A**). The CLAN incidence rate in unselected cells was somewhat higher, albeit still small, vs. that seen in the original GTM3L cells (0.04%)^22^, which is likely due to the use of GTM3-high GFP expression cells (described above)

**Figure 1.**
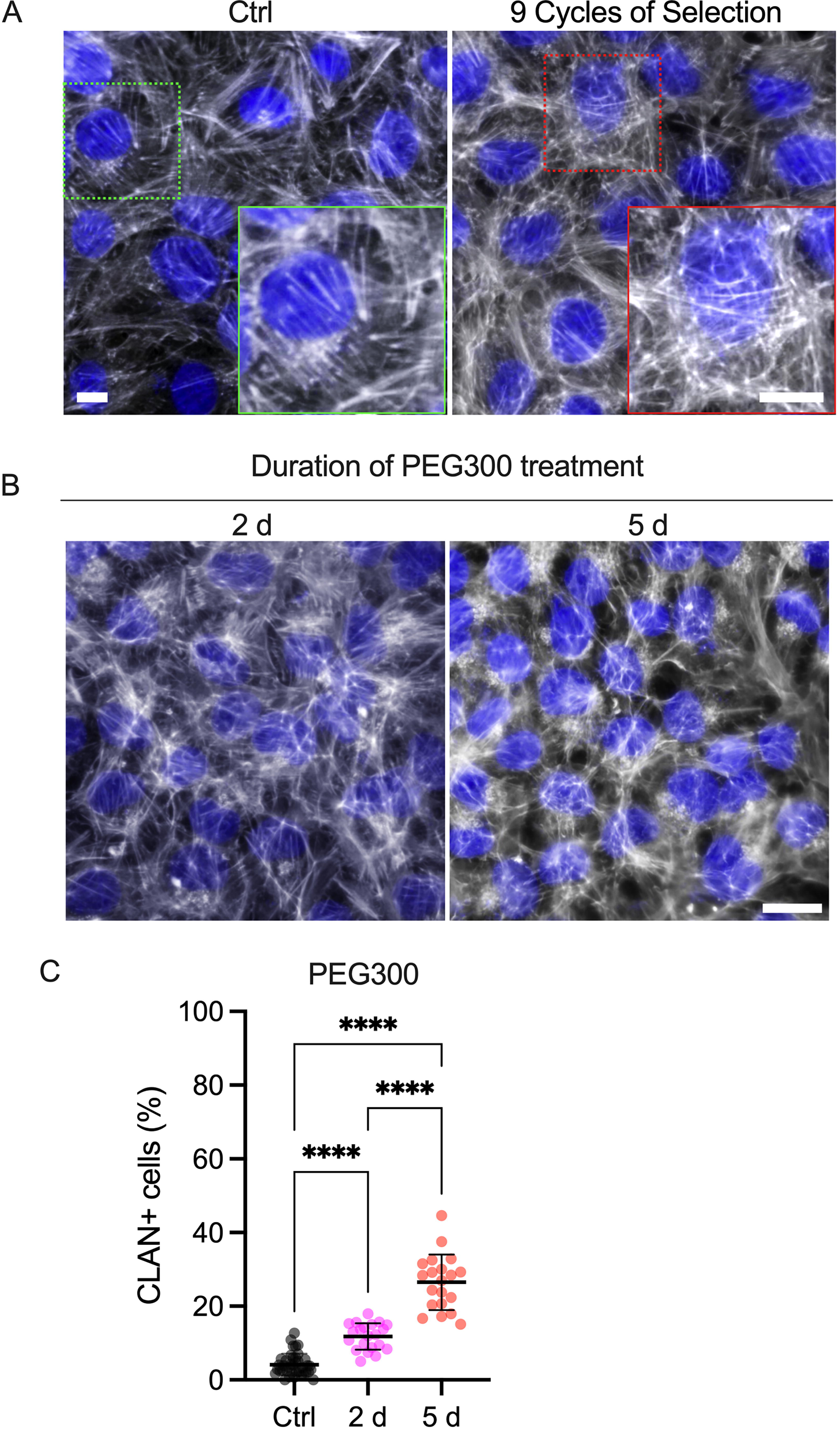
Enrichment of CLAN+ GTM3L cells using adhesion-based selection and PEG300 deswelling. (A) Representative fluorescence micrographs of F-actin in GTM3L cells before and after 9 cycles of adhesion-based selection, with nuclei and F-actin labelling shown in blue and grey, respectively. The green inset in A shows a zoomed-in view of F-actin that does not form a CLAN. The red inset in B shows a zoomed-in view of F-actin in a CLAN. Scale bars, 10 μm. **(B)** Representative fluorescence micrographs of F-actin in GTM3L cells that underwent 9 cycles of adhesion-based selection and were then subjected to PEG300 treatment for either 2 or 5 days. Nuclei and F-actin are labelled in blue and grey, respectively. Scale bar, 20 μm. **(C)** Analysis of the percentage of CLAN+ cells caused by treatment with PEG300 for 2 and 5 days (n = 40 images from 12 experimental replicates for the control group, n = 20 images/group from 6 experimental replicates for the groups treated with 2% PEG300). The bars and error bars indicate means ± standard deviations. Significance was determined by one-way ANOVA using multiple comparisons tests. ****: p<0.0001.

#### Enrichment of CLAN+ cells using PEG300

Studies have shown that macromolecular crowding induced by PEG promotes CLAN formation in an acellular actin filament solution modeling system.^39, 40^ We hypothesized that cell shrinkage induced by PEG would also facilitate CLAN formation in GTM3L cells. To investigate the effect of cellular crowding on CLAN formation in GTM3L cells, cells that had undergone 9 cycles of adhesion-based selection (see above) were treated with 2% PEG300 for either 2 or 5 days, after which we quantified the incidence rate of CLAN+ cells. As expected, GTM3L cells exposed to PEG300 shrank (**Figure 1A and B**). Additionally, there was a marked increase in the incidence of CLAN+ cells compared to the control group, i.e. vs. cells that had been enriched by selecting for adhesion but without PEG300 deswelling (**Figure 1B**). Specifically, the CLAN+ incidence rate was 4.16 ± 2.95% in control cells, which increased to 11.80 ± 3.58% and 26.50 ± 7.51% after 2- and 5-days of PEG300 treatment, respectively (**Figure 1B and C**).

We further studied whether withdrawal of PEG300 would lead to loss of CLANs. GTM3L cells were treated with PEG300 for 4-7 days to induce CLAN formation, followed by replacement of the PEG300-containing culture media with PEG-free media. We observed a time-dependent loss of CLANs after PEG300 withdrawal (**Figure 2A-H and supplemental video)**. Interestingly, while CLAN formation required several days, most of the PEG300-induced CLANs were lost within a few hours after PEG300 withdrawal (CLAN persistence = 58.94 ± 61.70 min [mean ± SD], N=31 cells; **Figure 2I**).

**Figure 2.**
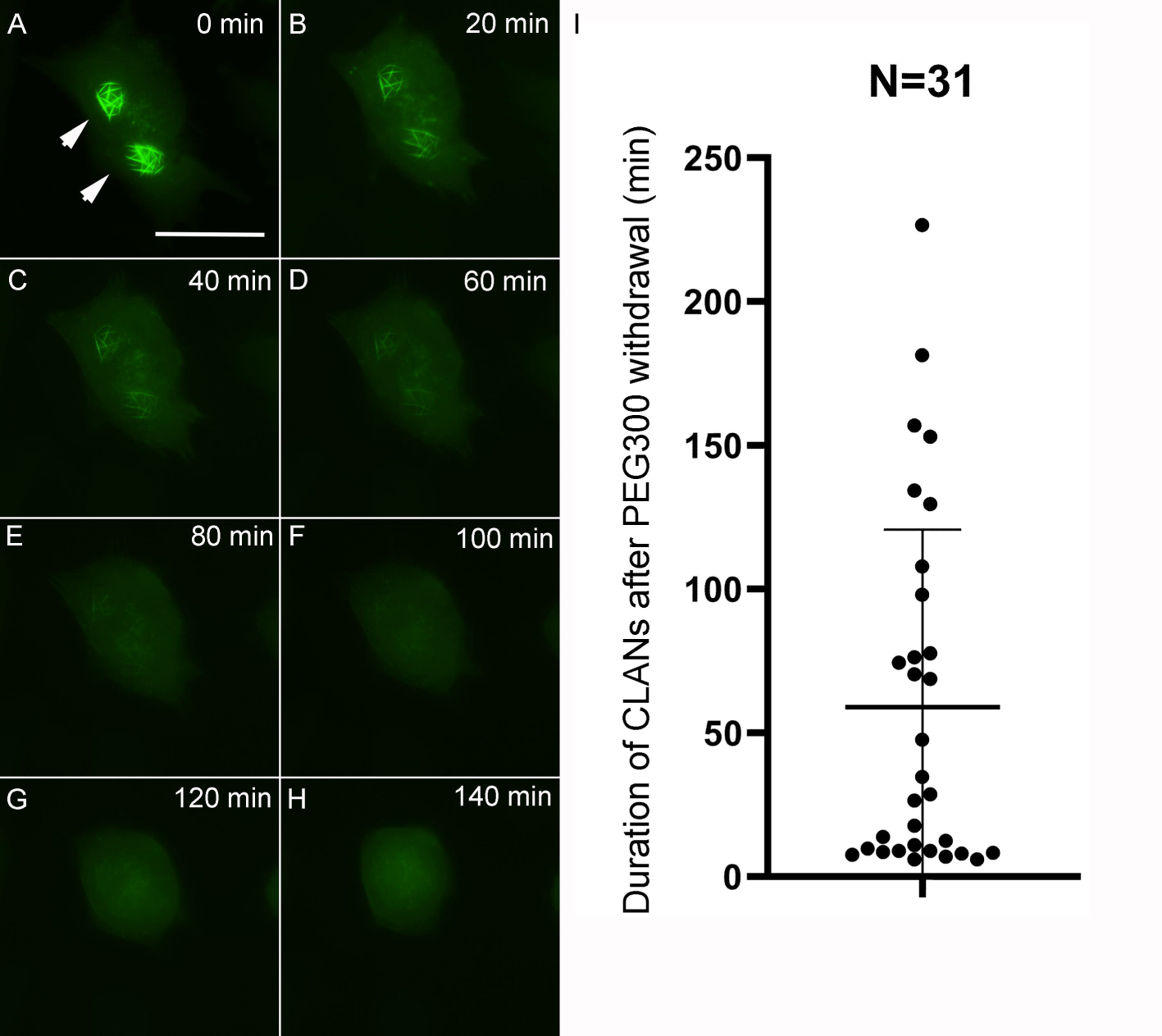
The persistence of CLANs after PEG300 withdrawal. (A-H) Images of a GTM3L cell at various times after PEG300 was removed from cells by medium change. Arrowheads denote CLANs in live cells. Scale bar: 50µm. **(I)** The persistence time of CLANs after PEG300 withdrawal, as assayed in in 31 GTM3L cells. Bars: mean and SD.

#### Enrichment of CLAN+ cells based on cell stiffness

We recently showed that GTM3L cells containing CLANs are stiffer than cells without CLANs.^22^ Therefore, we asked whether we could further enrich CLAN+ cells using a well-established microfluidic device that sorts cells depending on their stiffness.^27–31^ GTM3L cells, after undergoing 9 cycles of adhesion-based selection, were passed through this device, which contains a narrow channel with angled constrictions that direct cells along a stiffness-dependent trajectory, thereby sorting cells into four groups according to their stiffness. We note that no cells were obtained from one of the five outlets, and thus have only 4 groups which we denote as extra soft, soft, medium, and stiff. Note that these categorizations do not map onto cellular Young’s modulus values definable *a priori*, since the sorting process depends on cell stiffness, channel geometry, and cell size and shape.^27–31^ To evaluate the capability of these sorted cells to form CLANs, we then subjected them to a 5-day treatment with PEG300, using unsorted cells as a control. We established an association between the stiffness of GTM3L cells, based on the microfluidic-based cell sorting, and their ability to form CLANs. Specifically, we observed a significant elevation in CLAN formation as cell stiffness increased (**Figure 3A**).

**Figure 3.**
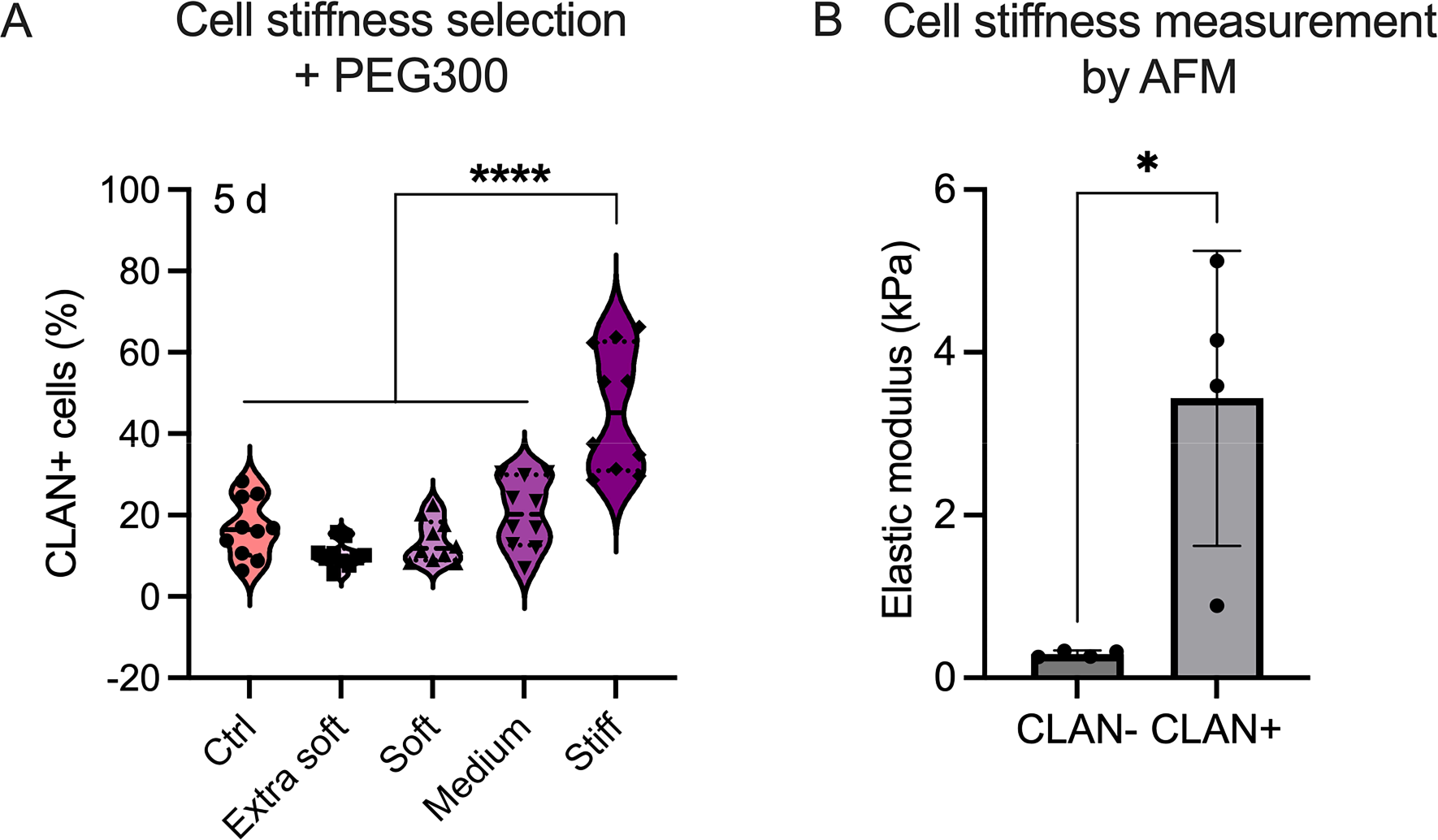
Enrichment of CLAN+ GTM3L cells based on cell stiffness. (A) GTM3L cells, after 9 cycles of adhesion-based selection, were sorted into four groups according to their stiffness: extra soft, soft, medium, and stiff. Control cells were unsorted cells. Each group of cells were plated on coverslips and exposed to 2% PEG300 for 5 days. The incidence rate of CLAN+ cells was quantified in each cell group (n = 10 images/group from 3 experimental replicates). The violin plot shows medians as horizontal solid lines and the interquartile range as dashed lines. Significance was determined by one-way ANOVA using multiple comparisons tests. ****: p<0.0001. **(B)** Stiffness of CLAN+ and CLAN-cells in an enriched GTM3L cell population was measured by AFM with a 10 μm tip (n = 4 cells/group). The bars and error bars indicate means ± standard deviations. Significance was determined by unpaired Student’s t-test. *: p<0.05.

Together, using a combination of all 3 CLAN+ cell enrichment methods (9 cycles of adhesion-based selection, cell sorting based on cell stiffness, and PEG300 treatment), we created a GTM3L sub-population containing about 50% CLAN+ cells (**Figure 3A**). This represents a significant enrichment from the original 0.28% CLAN+ incidence in GTM3L cells.

### Confirmation of cell stiffening in enriched GTM3L cells

Motivated by our previous findings that CLAN+ GTM3L cells are stiffer than CLAN-GTM3L cells and naïve GTM3 cells^22^, we also measured cell stiffness using AFM. We noted that in both our current and previous ^22^ studies, the CLAN+ GTM3L cells were reported to have a stiffness of approximately 4 kPa. Importantly, we confirmed that CLAN+ cells were stiffer than CLAN-cells (N=4 cells/group, p<0.05) (**Figure 3B**). This strengthens our confidence in our approach to selecting CLAN+ cells from a mixed population.

### Substrate stiffness influences CLAN incidence in GTM3L cells

Substrate stiffness has been shown to affect cell stiffness, with cells adapting by increasing their own stiffness in response to being cultured on a more rigid substrate.^41, 42^ In our study, we observed a notable correlation between the stiffness of cells and the presence of CLANs. Therefore, we hypothesized that substrate stiffness might also impact the formation of CLANs in GTM3L cells. To explore this hypothesis, we cultured stiff GTM3L cells, enriched through adhesion-based and cell stiffness selection steps (see above), on two different substrates: soft hydrogels with a stiffness of approximately 2.36 kPa, and stiff glass coverslips. The cells were then treated with PEG300 for 5 days. We observed that GTM3L cells cultured on the stiff substrate exhibited a significantly elevated incidence of CLAN+ cells (p<0.0001) (**Figure 4**).

**Figure 4.**
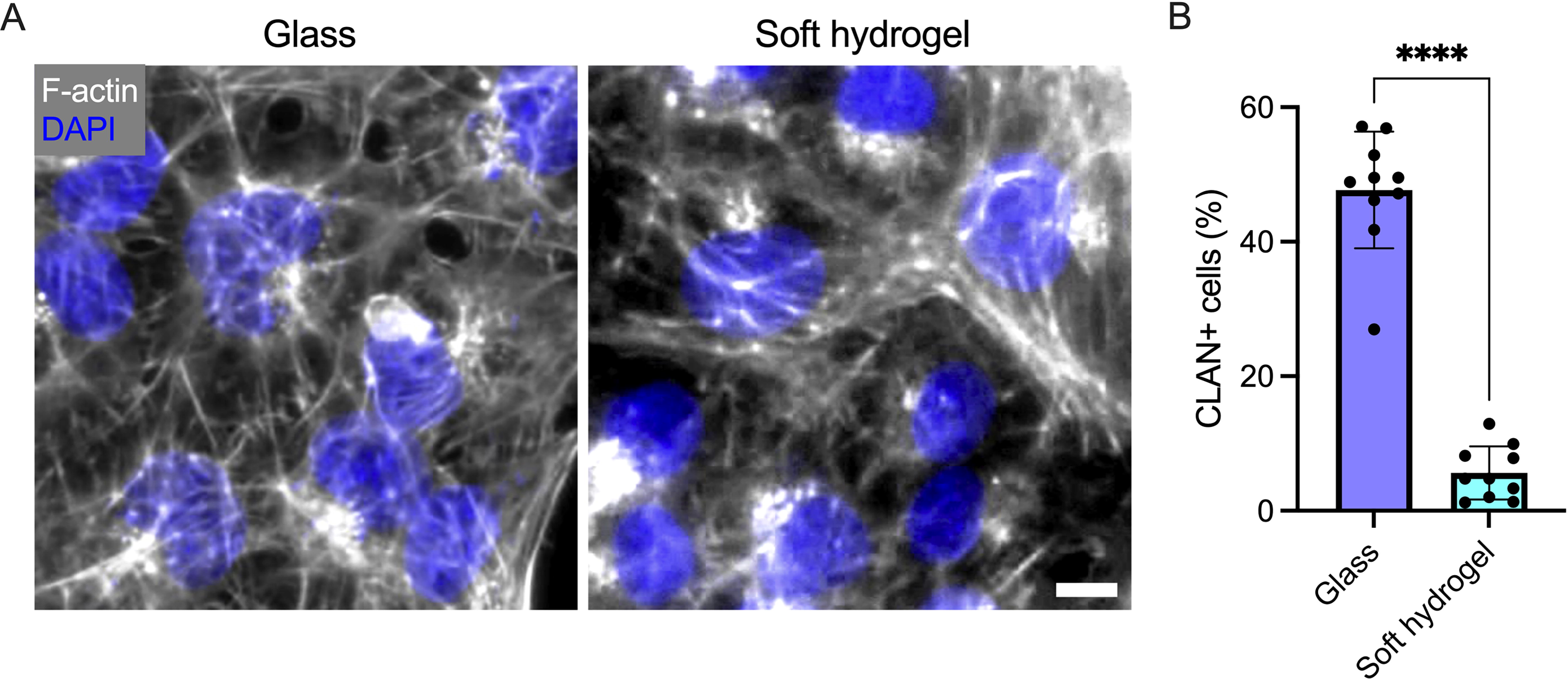
Substrate stiffness affects CLAN formation. (A) Representative fluorescence micrographs showing F-actin in GTM3L cells, with CLAN incidence rate enriched through adhesion- and cell stiffness-based selection. Cells were cultured on either glass coverslips or soft hydrogels (2.36 kPa). Nuclei are labelled in blue. Scale bar, 20 μm. **(B)** Incidence rate of CLAN+ cells as a function of substrate stiffness (n = 10 images/group from 3 experimental replicates). The bars and error bars indicate means ± standard deviations. Significance was determined by unpaired Student’s t-test. ****: p<0.0001.

### The mechanical properties of CLAN-like networks and CLAN-containing nucleus

To further study the impact of CLANs on cell biomechanics, we used agent-based models to probe the rheological properties of CLANs and the role of CLANs in cell stiffening. First, we created a CLAN-like network with a triangular lattice geometry (**Figure 5A**) and performed bulk rheology measurements imposing shear, tensile, and compressive strains on the network (**Figure 5B**). We found that network stiffness was greatest when delivering tensile strain and the smallest when delivering shear strain (**Figure 5B**), implying that CLANs in cells resist tensile deformation effectively. Next, we performed simulations with only a cell nucleus or with a cell nucleus surrounded by CLANs with or without physical links between the nucleus and CLANs (**Figure 5C and D**). When compressive strain was applied to the two types of structures, resistance to compression was higher when there were physical links between the nucleus and CLANs (**Figure 5C and D**). This implies that it is more difficult to deform the nucleus in the presence of CLANs, which is consistent with the increase in cell stiffness measured experimentally by AFM.

**Figure 5.**
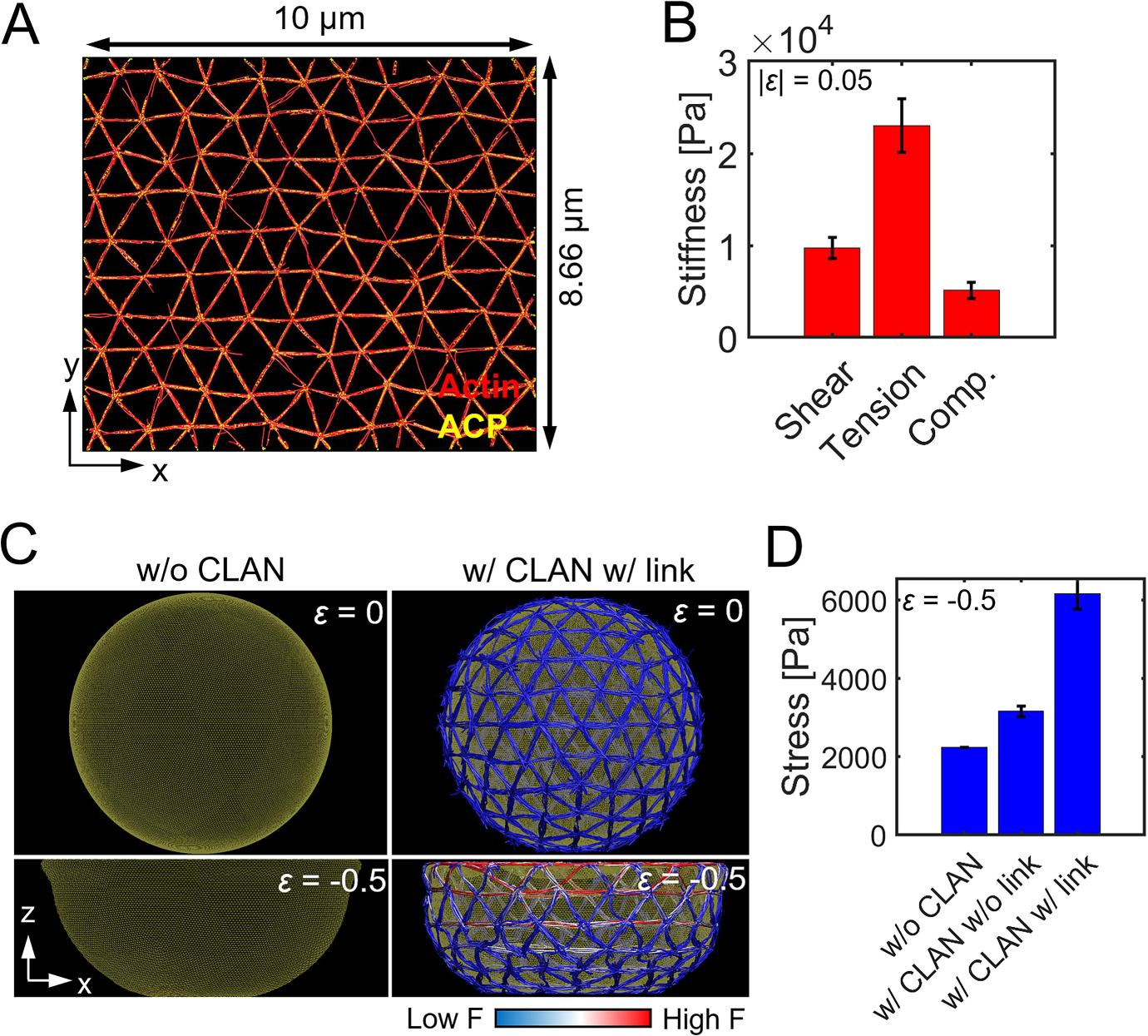
Simulations of the biomechanical effects of CLAN-like networks. (A) A CLAN-like network consisting of F-actin (red) and actin cross-linking proteins (ACPs, yellow) created in a thin rectangular domain. **(B)** Stiffness of the CLAN-like network in (A) in response to shear, tensile, and compressive deformations. The bars and error bars indicate means ± standard deviations (N=10). Significance was determined by one-way ANOVA and Tukey post-hoc test with p < 0.0001 by pairwise comparison between deformation types. **(C)** Snapshots of the nucleus with or without CLANs before and after the application of compressive normal strain up to ε = -0.5. In the case with CLANs, there are physical links between a fraction (20%) of F-actins and the nucleus. Colors in the actin fibers indicate the relative tensile force that fiber is bearing. Compressive forces are considered zero force and shown in blue. **(D)** Stress calculated from the resistant force developed during compressive deformations. The bars and error bars indicate means ± standard deviations (N=10). Significance was determined by one-way ANOVA and Tukey post-hoc test with p < 0.0001 by pairwise comparison between conditions.

## Discussion

In this study, we sought to understand various features of the enigmatic actin structures known as CLANs, which occur in TM cells and are associated with ocular hypertension. Towards this end, we used our previously reported GTM3L cells, which spontaneously form GFP-labelled CLANs in some cells. Fortunately, we were able to combine several methods to increase the incidence rate of CLAN+ GTM3L cells, successfully obtaining a subpopulation of GTM3L cells with ∼50% CLAN incidence rate. This significant enrichment of CLAN+ cells will be a valuable tool in future work, since it provides a large population of CLAN-positive cells to work with.

We also asked whether the amount of transgene expression in GTM3L cells was associated with the extent of lentiviral integration into the host cell’s genome. To our surprise, we found that all GTM3L cells, regardless of their transgene expression levels, had on average only one lentiviral insertion/integration event per cell. Generally, lentiviral integration sites in host chromosomes are random, and high dose of lentiviruses (more than 1 copy per cell) can insert at multiple regions in the host cell genome.^43, 44^

The above finding led us to the hypothesis that the presence or absence of CLANs in GTM3L cells depends on the locus of lentiviral insertion rather than the number of insertions. The rationale for this hypothesis includes the following:

- It is well known that lentiviruses integrate at random sites as well as at certain “hot spots” in the genome. Since the original GTM3 cells are monoclonal and all GTM3L cells were cultured in the same extracellular environment, the most likely explanation for CLAN formation is the difference in lentiviral integration sites.
- Different from the monoclonal GTM3 cells, our GTM3L cells are polyclonal, i.e. we did not conduct clonal selection or expansion after transduction. We observed a very low and unpredictable CLAN formation rate in GTM3L cells: sometimes there were many CLAN+ cells in a well, and sometimes there were no CLAN+ cells at all. This observation is consistent with our hypothesis and suggests that only a few lentiviral integration patterns will trigger CLAN formation.

Overall, the formation of CLANs in GTM3L cells may require two “hits”. We hypothesized that GTM3L cells with certain lentiviral insertion/integration pattern(s) are prone to form CLANs. When these cells grow in an unfavorable environment for CLAN formation, they form only a few CLANs (∼0.28%), yet if they grow in a favorable environment, such as on a stiff substrate or in the presence of PEG300, they form significantly more CLANs.

We were able to enrich the population of CLAN+ cells using different methods: differential substrate adhesion, PEG300 exposure and cell stiffness-based sorting. Interestingly, these methods were synergistic, suggesting that they might be selecting for different CLAN-associated phenotypes. The timescale associated with PEG300 exposure was noteworthy, since the induction of CLANs by PEG300 took several days, yet CLAN disassembly after PEG300 removal occurred within hours. We do not understand the cause of this difference, nor do we understand the exact mechanism by which PEG300 induces CLANs in TM cells. The simplest possibility is that cell deswelling by PEG300 leads to cytoplasmic molecular crowding, an effect that was seen in an acellular model where crowding decreased α-actinin binding to F-actin and possibly led to F-actin thinning and shortening.^40^ Another possibility is that PEG300 exposure leads to cellular stress, and that CLAN formation is a generic stress response of TM cells. For example, the formation of CLANs at the perinuclear region might provide TM cells with additional mechanical shielding or stabilization. This idea that CLANs as a stress response may also help explain the observation that CLANs tend to form more on stiffer substrates, as well as in the presence of DEX and TGF-β2. More experiments are clearly needed to explore this concept.

We employed an agent-based computational model to understand why CLAN+ cells are stiffer. We hypothesized that CLAN+ cells exhibit higher stiffness because their nuclei are harder to deform due to the surrounding CLANs. To test this hypothesis, we first computationally measured the rheological properties of CLAN-like networks and found that the networks resisted tensile deformation effectively. Then, we applied compressive deformation to a simplified cell nucleus with or without CLANs and found that the presence of CLANs resulted in much higher resistance of the cell nucleus to compression due to high tensile resistance of CLANs; specifically, deformation of the cell nucleus was hindered by the limited extensibility of actin fibers. Based on these observations, which are consistent with our hypothesis, it is likely that the higher stiffness of CLAN+ cells partially originates from CLANs around the nucleus.

Probably the most important question about CLANs is whether they directly contribute to decreased outflow facility and hence elevated IOP, or are simply an associated epiphenomenon. Our data do not directly answer this question; even the finding that a stiff substrate, such as seen in the TM of ocular hypertensive eyes, promotes CLAN formation is consistent with both possibilities. To address this question will require more work in a variety of models, and we suggest that our findings can play an important role by motivating and facilitating cell-based models. For example, one possibility would be to decellularize TM tissue in perfused anterior segments and then repopulate the TM by magnetically steering CLAN+ cells into the TM. Another approach would be to use an artificial outflow pathway construct (TM-on-a-chip), populating the construct with CLAN+ cells to determine effects on flow resistance. All such studies will require large numbers of CLAN+ cells, the production of which will be greatly facilitated by our enrichment strategies. Also, if the gene(s) that promotes CLAN formation are discovered (e.g. by comparing CLAN- vs. CLAN+ GTM3L cells using DNAseq), CLAN formation can be induced in the mouse TM and the effect of CLANs on outflow facility and IOP can be determined in vivo.

Of course, this work is subject to certain limitations. Besides the lack of outflow facility and IOP data, key among these is that the GTM3L cells are derived from a transformed cell line (GTM cells). It is well known that the biology of transformed TM cells is different from primary TM cells.^21^ However, we believe these GTM3L are still a valuable tool for studying CLANs because of several reasons.

1. Important features are consistent between GTM3L CLAN+ cells and pHTM CLAN+ cells, such as resistance to actin relaxing reagents^16, 22^ and increased cell stiffness.^22^
2. Our GTM3L cells form CLANs spontaneously, offering several advantages:

• Live imaging of CLAN+ cells under physiological conditions.
• Enriched CLAN+ GTM3L cell populations make omics-based studies possible.
Our GTM3L cell line has unlimited proliferation capability, and CLAN formation in this line is spatially consistent (predominantly in the perinuclear region), which improves reproducibility.
3. LifeAct-GFP expression does not affect TM cell biomechanical properties. Unlike human mesenchymal stem cells,^45^ our GTM3L cells did not demonstrate adverse effects since CLAN-GTM3L cells and GTM3 cells showed similar stiffness and viscosity.
4. Our cell culture studies are directly intellectually linked to functional outcomes in whole eyes.

Specifically, we know that increased TM tissue stiffness is associated with OHT^5, 6^ and impaired IOP homeostasis, which in view of the greater stiffness of GTM3L CLAN+ cells, strongly suggests mechanistic link(s) between CLANs and OHT. In future, we fully expect to be able to translate our findings from GTM3L cells into whole eyes.

In summary, we have developed an effective strategy to greatly increase the presence of CLAN+ cells in our newly discovered GTM3L subline. Based on our findings, we believe that CLANs, either induced by glaucomatous signals (elevated TGFβ2, elevated cortisone or steroid treatment, or elevated IOP/mechanical stretching) or even as a primary initiating factor of glaucoma, lead to pathological changes in TM cell biology, biomechanics, and mechanobiology, resulting in OHT in glaucomatous eyes. Further research is needed to determine these changes and the underlying mechanisms, and this subline will be a useful tool for this purpose.

## Acknowledgements

Supported by the National Institute of Health/National Eye Institute Award Numbers R01EY026962 (WM), R01EY031700 (WM), R21EY033929 (WM) and R01EY031710 (CRE);

BrightFocus Foundation G2023009S (WM); a challenge grant from Research to Prevent Blindness (Department of Ophthalmology, Indiana University School of Medicine); the Georgia Research Alliance (CRE); as well as National Science Foundation Award Number 2134701 (TS) and CBET-2225476 (TS).

The authors thank Dr. Kenneth Cornetta of the Gene Therapy Testing Laboratory at Indiana University School of Medicine for ddPCR resources.

The content is solely the responsibility of the authors and does not necessarily represent the official views of the National Institutes of Health.

