## Supplemental materials for "Characterization, enrichment, and computational modeling of cross-linked actin networks in trabecular meshwork cells"

**Droplet digital PCR conditions**

**Primer	Sequence	Final Concentration [nM]**

ApoB-F	5’-tgaaggtggaggacattcctcta-3’	900

ApoB-R	5’-ctggaattgcgatttctggtaa-3’	900

TP-ApoB	5’-HEX-cgagaatcaccctgccagacttccgt	250

WPRE-F	5’-CACCACCTGTCAGCTCCTTT–3’	900

WPRE-R	5’-ACAACACCACGGAATTGTCA-3’	900

TP_-WPRE	5’-FAM-ACTTTCGCTTTCCCCCTCCCTATT	250

**PCR condition	Number of Cycles	Settings**

Step 1	1	Hold at 95˚C for 10 minutes

Step 2	40	94˚C for 30 seconds

60˚C for 1 minute

Step 3	1	Hold at 98˚C for 10 minutes

Step 4	∞	1	Hold at 12˚C

**Equipment / Reagent	Supplier / Model / Cat #**

Droplet Reader	Bio-Rad / QX200

Droplet Generator	Bio-Rad / QX200

Plate Sealer	Bio-Rad / PX1

Thermocycler	Bio-Rad / C1000 Touch

DG8 cartridges	Bio-Rad / 1864008

DG8 Gaskets	Bio-Rad / 1863009

ddPCR 96 well plates	Bio-Rad / 12001925

Foil plate seals	Bio-Rad / 1814040

Droplet Generation Oil	Bio-Rad/1863005

Water	HyClone/SH30538

ddPCR SuperMix for Probe (No dUTP), 2X	Bio-Rad / 1863024

Restriction Enzyme HindIII [20 U/uL]	NEB / R3104S

Droplet reader oil	Bio-Rad / 1863004

**Sorting of GTM3L cells by GFP intensity**

Cultured GTM3L cells were suspended using trypLE and sorted based on GFP intensity (FITC-A channel), as shown in Supplemental Figure 1.

**Agent-based model**

*Brownian dynamics simulations via the Langevin equation*

In the model, all elements are defined by points (i.e., nodes). The displacements of points are governed by the Langevin equation with inertia neglected:

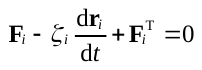
(S1)

where **r***_i_* is the position of the *i*th point, *ζ_i_* is a drag coefficient, *t* is time, **F***_i_* is a deterministic force, and **F**^T^*_i_* is a stochastic force. The magnitude **F**^T^*_i_* of is determined by the fluctuation-dissipation theorem:

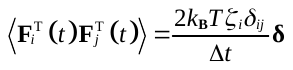
(S2)

where *δ_ij_* is the Kronecker delta, **δ** is a second-order tensor, *k*_B_*T* is thermal energy, and Δ*t* is a time step. The drag coefficients of points for cylindrical segments are computed using an approximated form ^1^:

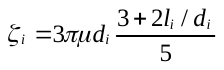
(S3)

where *μ* is medium viscosity, and *l_i_* and *d_i_* are the length and diameter of a cylindrical segment connected to the point, respectively. The drag coefficient of nodes for a nuclear membrane is determined based on membrane thickness and a distance between adjacent nodes. Using the Langevin equation, the velocity of each point, d**r***_i_*/d*t*, is calculated. The positions of all points are updated in each time step using the Euler integration scheme and the calculated velocities:

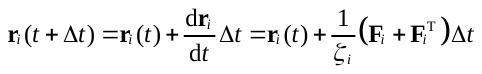
(S4)

Deterministic forces include extensional forces that maintain the equilibrium lengths of cylindrical segments or membrane chains, bending forces which maintain angles formed by interconnected segments or dihedral angles on the membrane close to their equilibrium values, and repulsive forces that represent volume-exclusion effects acting between overlapping components. The extensional and bending forces originate from the following harmonic potentials:

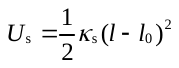
 (S5)
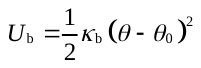
(S6)

where *κ*_s_ is extensional stiffness, *l* and *l*_0_ are instantaneous and equilibrium lengths, *κ*_b_ is bending stiffness, and *θ* and *θ*_0_ are instantaneous and equilibrium angles, respectively.

*Simplification and mechanics of cytoskeletal components*

F-actin is simplified into serially connected cylindrical segments. Each segment has polarity defined by barbed and pointed ends (Supplemental Figure 2A). Extensional (*κ*_s,A_) and bending (*κ*_b,A_) stiffnesses of F-actin regulate the equilibrium length of actin segments (*l*_0,A_ = 140 nm) and an equilibrium angle formed by two adjacent actin segments (*θ*_0,A_ = 0 rad), respectively. The persistence length of F-actin (*l*_p_) with *κ*_b,A_ is ~9 μm.^2^

Actin cross-linking proteins (ACPs) consist of two cylindrical segments connected at their center point. Extensional (*κ*_s,ACP_) and bending (*κ*_b,ACP_) stiffnesses of ACPs maintain the equilibrium length of each ACP segment (*l*_0,ACP_ = 23.5 nm) and an equilibrium angle formed by two ACP segments at the center point (*θ*_0,ACP_ = 0 rad), respectively.

Forces exerted on actin segments by bound ACPs are distributed to the barbed and pointed ends of the actin segments as described in our previous work.^3^ In actin structures, a repulsive force is calculated only between F-actins, following a harmonic potential:^4^

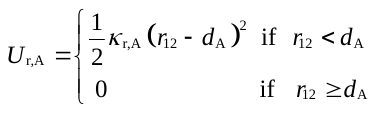
(S7)

where *κ*_r,A_ is the strength of the repulsive force, *r*_12_ is a minimum distance between two neighboring actin segments, and *d*_A_ is the diameter of the actin segments.

*Simplification and mechanics of the nucleus*

A cell nucleus is simplified into a spherical triangulated mesh with the initial radius of 2.5 µm and with area and volume conservation (Supplemental Figure 2C). Extensional stiffness of the nucleus (*κ*_s,N_) governs the equilibrium length of chains between mesh nodes (*l*_0,N_ = ~70 nm). Bending stiffness of the nucleus (*κ*_b,N_) regulates dihedral angles formed by adjacent triangular elements on the mesh close to their equilibrium value (*θ*_0,N_ = 0 rad). The area of each triangular element and the entire volume defined by the nucleus are conserved via the following harmonic potentials:

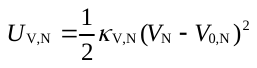
(S8)

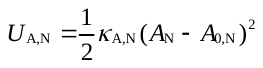
 (S9)

where *κ*_V,N_ and *κ*_A,N_ are the strengths of volume and area conservation, *V*_N_ is volume, *A*_N_ is area, and the subscript 0 denotes the equilibrium value.

A repulsive force is applied between triangular elements on the mesh with strength, *κ*_r,N_, to prevent them from crossing each other. The repulsive force originates from the following potential:

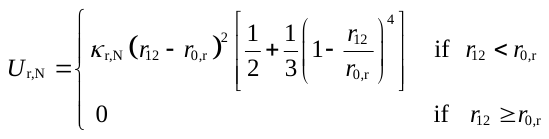
(S10)

where *κ*_r,N_ is the strength of the repulsive force, and *r*_0,r_ is a distance below which the repulsive force starts acting. In this case, *r*_0,r_ is equal to the thickness of the nucleus mesh (*r*_0,r_ = *d*_N_). Note that this repulsive force increases more rapidly as the two nucleus elements get closer to each other.

*Actin dynamics*

The formation of F-actin is initiated from a nucleation event occurring with a constant rate constant, *k*_n,A_, with the appearance of one cylindrical segment. The polymerization of F-actin is simulated by adding cylindrical segments at the barbed or pointed end of existing filaments with a constant rate constant, *k*_+,A_. The nucleation and polymerization events can take place only if a new segment emerging from the events is located within a designated space for network assembly. The depolymerization of F-actin does not occur.

In simulations only with a network, the nodes of a two-dimensional triangular lattice whose node-to-node distance is ~1 µm are used as locations for the actin nucleation event. Noise is applied to the nucleation locations to prevent the network from being a perfect triangular lattice. The orientation of new segments emerging from the nucleation event is determined by one of the six unit directional vectors directed toward adjacent nodes. The initial segment with 140 nm in length can elongate by up to 840 nm in the barbed-end direction and by up to 280 nm in the pointed-end direction, meaning that the maximum length of F-actin is 1.26 µm. Part of F-actins polymerized from the barbed end are connected to F-actins polymerized from adjacent nodes by ACPs to form small bundles between nodes (Supplemental Figure 2B). Part of F-actins polymerized from the pointed end are connected to other F-actins polymerized from the same node by ACPs to form connections between F-actins on each node.

In simulations with a nucleus, a CLAN-like network is created on the nucleus surface. For creating the CLAN-like network, the nodes of a triangular lattice defined along the nucleus surface are used as positions for the actin nucleation event. The node-to-node distance is ~1 µm. The way to create the network on the nucleus surface is the same as that used for the two-dimensional network.

*Dynamic behaviors of ACPs*

ACP arms bind to binding sites located every 7 nm on actin segments with a constant rate constant, *k*_+,ACP_, without preference for cross-linking angle. They can also unbind from F-actin at a force-dependent rate determined by Bell’s law:^5^

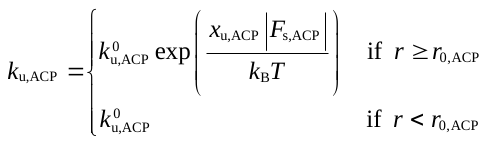
(S11)

where
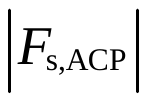
 is a spring force acting on an ACP arm, *k*^0^_u,ACP_ is the zero-force unbinding rate constant, and *x*_u,ACP_ is sensitivity to the applied force.

*Interactions between the cytoskeletal elements and the nucleus*

Repulsive forces are applied between all cytoskeletal elements and the nucleus mesh to prevent the cytoskeletal elements from entering the inside of the nucleus. The repulsive forces originate from the potential shown in Eq. S10 with the same strength, *κ*_r,N_. *r*_0,r_ is equal to the average of nuclear membrane thickness and the diameter of a neighboring element, *r*_0,r_ = (*d*_N_ + *d_i_*)/2, where *i* is either A (actin) or ACP.

A fraction of the endpoints of actin segments feel an attractive force from the nucleus if a distance between them and the nucleus is smaller than *d*_A_ + *d*_N_ to mimic physical links formed by the LINC complex ^6^. The fraction is set to 0 or 0.2. The attractive force is defined by the following harmonic potential:

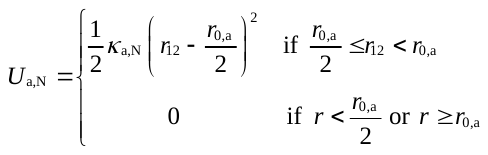
(S12)

where *κ*_a,N_ is the strength of the attractive force, and *r*_0,a_ is equal to *d*_A_ + *d*_N_**.** Note that the attractive force is always applied in a direction normal to the closest nucleus triangular element, meaning that the endpoints can slide along the nucleus surface with the maintenance of the equilibrium distance from the nucleus. Drag forces acting between the endpoints and the nucleus are ignored for simplicity, meaning that the endpoints bound to the nucleus still experience a drag force defined by Eq. S3.

**Supplemental Figures**

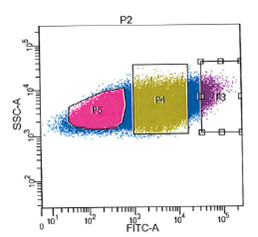

**Supplemental Figure 1. Isolation of low-, medium-, and high-GFP expressing GTM3L cells using fluorescence-activated cell sorting.** Abbreviations: SSC: side scattering. P5: low GFP; P4: medium GFP; P3: high GFP.

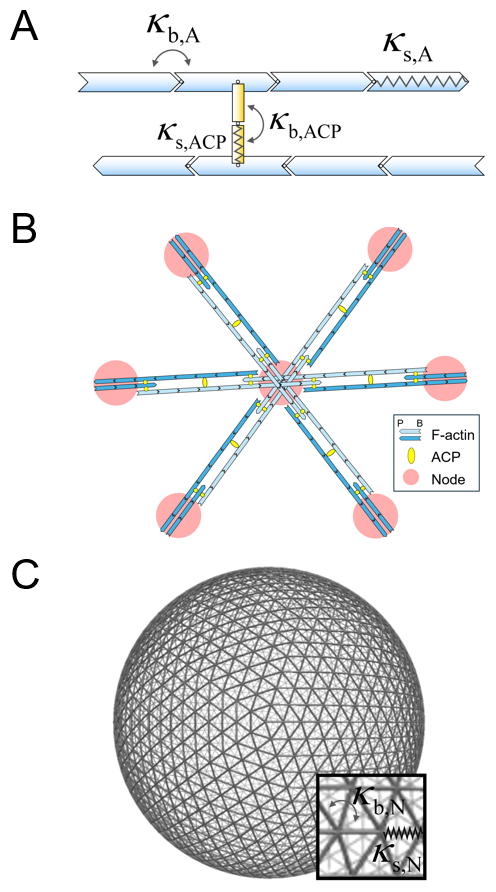

**Supplemental Figure 2. Agent-based model details.** **(A)** A CLAN-like network consists of F-actin (cyan) and actin cross-linking proteins (ACP; yellow). Extensional (*κ*_s_) and bending (*κ*_b_) stiffnesses maintain the equilibrium length of segments and the equilibrium angle formed by segments, respectively. F-actin has polarity defined by barbed and pointed ends. **(B)** The CLAN-like network is created by the nucleation and polymerization of F-actin from nodes (pink) and the formation of cross-linking points between F-actins. Cyan indicates F-actins forming from the center node, and dark blue indicates F-actins elongating from six adjacent nodes. **(C)** A cell nucleus is simplified into a triangulated mesh with the conservation of volume and surface area. Extensional (*κ*_s,N_) and bending (*κ*_b,N_) stiffnesses maintain the equilibrium length of chains between nodes and the equilibrium angle formed by adjacent chains, respectively.

**Supplemental Table**

| **Symbol** | **Definition** | **Value** |
| --- | --- | --- |
| *l*_0,A_ | Length of an actin segment | 1.4×10^-7^ [m] |
| *d*_A_ | Diameter of an actin segment | 7.0×10^-9^ [m] ^7^ |
| *θ*_0,A_ | Bending angle formed by adjacent actin segments | 0 [rad] |
| *κ*_s,A_ | Extensional stiffness of F-actin | 1.69×10^-2^ [N/m] |
| *κ*_b,A_ | Bending stiffness of F-actin | 2.64×10^-19^ [N·m] ^2^ |
| *κ*_r,A_ | Strength of repulsive force b/w F-actins | 1.69×10^-3^ [N/m] |
| *k*_n,A_ | Nucleation rate of actin | 1.0×10^-4^ [μM^-1^s^-1^] |
| *k*_+,A_ | Polymerization rate of actin at the barbed end | 60 [μM^-1^s^-1^] |
| *l*_0,ACP_ | Length of an ACP arm | 2×10^-8^ [m] ^8^ |
| *d*_ACP_ | Diameter of an ACP arm | 1.0×10^-8^ [m] |
| *θ*_0,ACP_ | Bending angle formed by two ACP arms | 0 [rad] |
| *κ*_s,ACP_ | Extensional stiffness of ACP | 2.0×10^-3^ [N/m] |
| *κ*_b,ACP_ | Bending stiffness of ACP | 1.04×10^-19^ [N·m] |
| *k*_+,ACP_ | Binding rate of an ACP arm | 100 [s^-1^] |
| 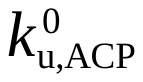 | Zero-force unbinding rate constant of ACP | 0.01 [s^-1^] |
| *x*_u,ACP_ | Sensitivity of ACP unbinding to applied force | 1.0×10^-10^ [m] |
| *l*_0,N_ | Length of a nucleus mesh chain | 7.0×10^-8^ [m] |
| *d*_N_ | Thickness of a nucleus mesh | 5.0×10^-8^ [m] |
| *θ*_0,N_ | Dihedral angle formed by two adjacent mesh elements | 0 [rad] |
| *κ*_s,N_ | Extensional stiffness of a nucleus mesh chain | 7.2×10^-4^ [N/m] |
| *κ*_b,N_ | Bending stiffness of a nucleus mesh | 6.45×10^-19^ [N·m] |
| *A*_0_,_N_ | Area of one triangular mesh | 2.12×10^-15^ [m^2^] |
| *κ*_A,N_ | Strength of areal conservation | 1.0×10^-5^ [N/m] |
| *V*_0_,_N_ | Volume within a nucleus mesh | 6.54×10^-17^ [m^3^] |
| *κ*_V,N_ | Strength of volume conservation | 1.0×10^3^ [N/m^2^] |
| *κ*_a,N_ | Strength of attractive force b/w a mesh and ACPs | 1.0×10^-3^ [N/m] |
| *κ*_r,N_ | Strength of repulsive force b/w all cytoskeletal elements and a mesh and b/w nucleus meshes | 5.1×10^-3^ [N/m] |
| Δ*t* | Time step | 1.15×10^-4^ [s] |
| *μ* | Viscosity of a medium | 8.6 [Pa·s] |
| *k*_B_*T* | Thermal energy | 4.142×10^-21^ [J] |

**Supplemental Table 1.** **List of parameters employed in the model and their values.**
